# Prolonged exposure to lung-derived cytokines is associated with inflammatory activation of microglia in patients with COVID-19

**DOI:** 10.1101/2023.07.28.550765

**Authors:** Rogan A. Grant, Taylor A. Poor, Lango Sichizya, Estefani Diaz, Joseph I. Bailey, Sahil Soni, Karolina J. Senkow, Xochítl G. Pérez-Leonor, Hiam Abdala-Valencia, Ziyan Lu, Helen K. Donnelly, Robert M. Tighe, Jon W. Lomasney, Richard G. Wunderink, Benjamin D. Singer, Alexander V. Misharin, GR Scott Budinger, The NU SCRIPT Investigators

**Affiliations:** Division of Pulmonary and Critical Care Medicine, Department of Medicine, Feinberg School of Medicine, Northwestern University, Chicago, IL, USA; Division of Pulmonary, Allergy, and Critical Care Medicine, Duke University School of Medicine, Duke University, Durham, NC, USA; Department of Pathology, Feinberg School of Medicine, Northwestern University, Chicago, IL, USA; Department of Biochemistry and Molecular Genetics, Department of Medicine, Feinberg School of Medicine, Northwestern University, Chicago, IL, USA

## Abstract

Neurological impairment is the most common finding in patients with post-acute sequelae of COVID-19. Furthermore, survivors of pneumonia from any cause have an elevated risk of dementia^1–4^. Dysfunction in microglia, the primary immune cell in the brain, has been linked to cognitive impairment in murine models of dementia and in humans^5^. Here, we report a transcriptional response in human microglia collected from patients who died following COVID-19 suggestive of their activation by TNF-ɑ and other circulating pro-inflammatory cytokines. Consistent with these findings, the levels of 55 alveolar and plasma cytokines were elevated in a cohort of 341 patients with respiratory failure, including 93 unvaccinated patients with COVID-19 and 203 patients with other causes of pneumonia. While peak levels of pro-inflammatory cytokines were similar in patients with pneumonia irrespective of etiology, cumulative cytokine exposure was higher in patients with COVID-19. Corticosteroid treatment, which has been shown to be beneficial in patients with COVID-19^6^, was associated with lower levels of CXCL10, CCL8, and CCL2—molecules that sustain inflammatory circuits between alveolar macrophages harboring SARS-CoV-2 and activated T cells^7^. These findings suggest that corticosteroids may break this cycle and decrease systemic exposure to lung-derived cytokines and inflammatory activation of microglia in patients with COVID-19.

---

Nearly 800 million people have been diagnosed with Coronavirus Disease 2019 (COVID-19) and 7 million people have died. In the US alone, there are currently more than 100 million survivors of COVID-19^8^. Symptoms of post-acute sequelae of COVID-19 (PASC) are dominated by neurological, cognitive, and psychiatric dysfunction^4,9–11^. Cognitive impairment appears to be particularly common and long-lasting. A meta-analysis of world health records estimated that 2% of all symptomatic SARS-CoV-2 infections result in at least short-term cognitive impairment, with more than 50% of patients presenting to PASC centers reporting psychiatric symptoms and 15% of patients with PASC reporting sustained symptoms one year after initial infection^12^. An increased risk of dementia persists for at least two years after severe COVID-19^4^, similar to what is observed following pneumonia secondary to other pathogens^1–3^. These complications are more common in patients requiring ICU admission^13,14^. Mechanisms underlying cognitive impairment after recovery from COVID-19, however, are unknown.

## COVID-19 is associated with a transcriptional signature in microglia suggestive of NF-κB activation and cell-cycle arrest

We hypothesized that COVID-19 results in activation of microglia, an abundant and dynamic resident macrophage population in the central nervous system (CNS). This hypothesis is informed by the association between microglial activation and dementia in animal models and humans^5,15,16^. Accordingly, we collected frontal lobe samples at autopsy from five patients who died following SARS-CoV-2 infection and 3 patients who died without respiratory failure or sepsis between March, 2021 and April, 2022. Clinical features of these patients are included in Suppl. Table S1. We generated single-cell suspensions of these tissues and enriched them for live microglia, T cells, and other neuroimmune cells using flow cytometry sorting (Suppl. Fig. S1a). We then performed single-cell RNA-sequencing (scRNA-seq) on these samples, identifying 65,767 cell passing quality control, predominantly heterogeneous populations of microglia and CD8^+^ T cells (Fig. 1a-c).

**Figure 1.**
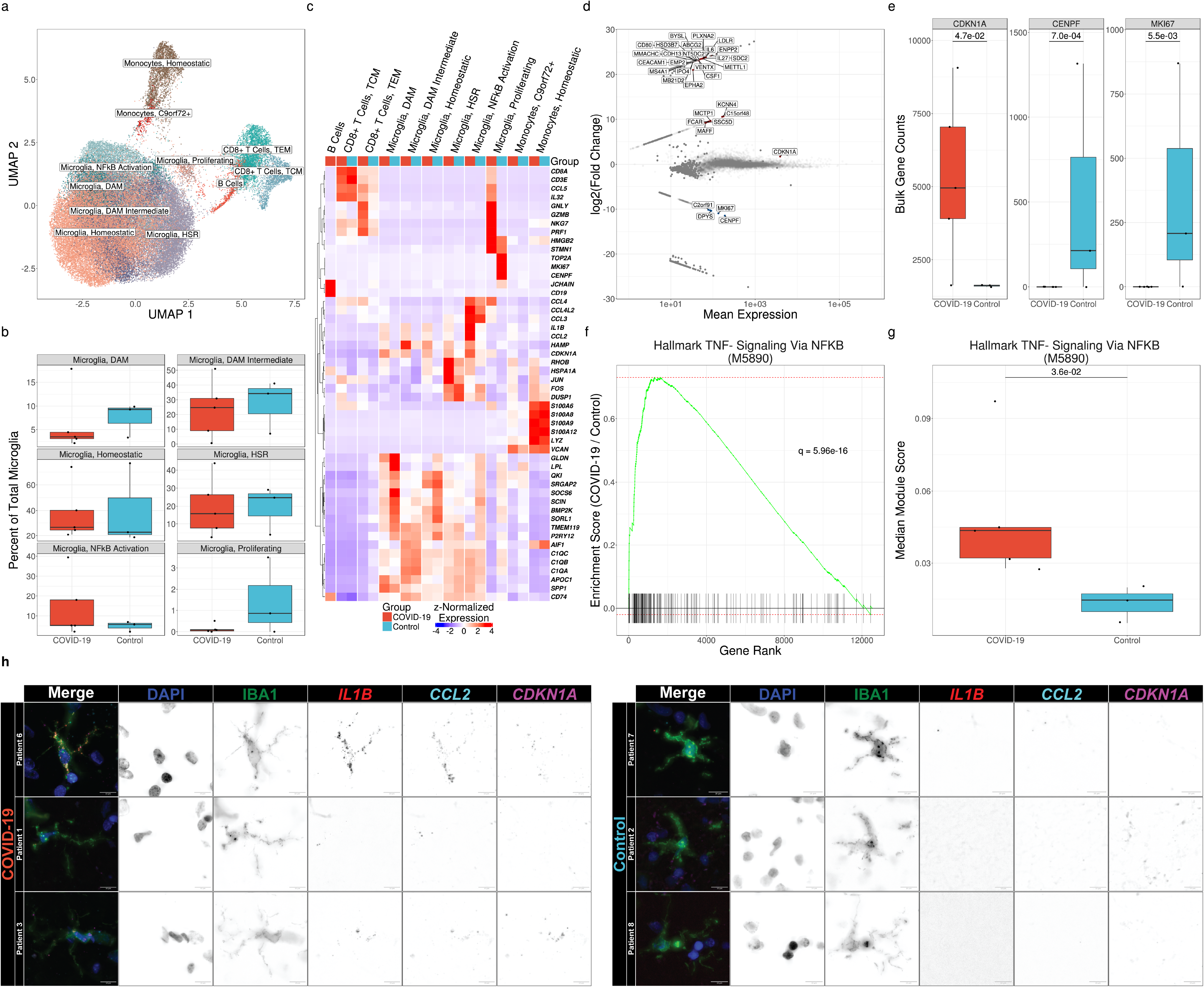
Microglia exhibit distinct transcriptional responses in patients with COVID-19. **(a)** UMAP of 65,767 cells isolated from the frontal lobes of 8 patients postmortem. **(b)** Relative abundance of microglial cell states as a percentage of total microglia. No significant differences are observed by diagnosis (q ≤ 0.05, Mann-Whitney). **(c)** Hierarchical clustering of mean marker gene expression by cell type and cell state by diagnosis. **(d)** MA plot of differentially expressed genes in total microglia in COVID-19 vs controls by pseudo-bulk differential expression analysis. Significantly upregulated genes are shown in red, and significantly downregulated genes are shown in blue (q < 0.05, Wald test). Genes shown in gray are not significantly differentially expressed. **(e)** Callouts of key markers of cell division and cell-cycle arrest from 1d. All genes shown are significantly differentially expressed (q < 0.05, Wald test). **(f)** Gene-set enrichment of Hallmark TNF-α Signaling Via NF-κB (M5890) from pseudo-bulk differential expression analysis (q = 5.96×10^-16^, multilevel splitting Monte Carlo). **(g)** Median modular expression of Hallmark TNF-α Signaling Via NF-κB (M5890) by diagnosis. Points represent median expression in total microglia from each patient (q = 2.6×10^-2^, Mann-Whitney). (**h)** Representative images of combined immunofluorescence and smFISH (RNAScope^TM^) from human frontal lobe tissue sections showing cell cycle-arrested, pro-inflammatory microglia in patients with COVID-19 relative to controls. Images are pseudo-colored by channel as follows. DAPI: blue, IBA1: green, *IL1B*: red, *CCL2*: cyan, *CDKN1A*: magenta.

Some investigators have suggested cognitive decline results from direct infection of the CNS by SARS-CoV-2. However, studies using robust techniques such as single-cell and single-nucleus RNA-seq and single-molecule fluorescence *in situ* hybridization (smFISH) have failed to identify consistent evidence of infection outside the lung^17–20^. Accordingly, we did not detect a single read of any SARS-CoV-2 gene or the negative sense genome scaffold required for replication^21^ when we aligned our scRNA-seq reads to a hybrid genome containing the human GRCh38.93 genome build and the wild-type Wuhan SARS-CoV-2 strain (NC045512.2) (Suppl. Fig. S1b). Investigators have also suggested that PASC may result from Epstein-Barr virus reactivation^22^. However, we did not detect a single read of any EBV1 gene when we included a linearized EBV1 genome (NC007605.1) in our hybrid genome (Suppl. Fig. S1c).

The development and progression of dementia has been associated with the accumulation of microglia with a distinct transcriptional phenotype defined by the expression of genes including *Apoe*, *Spp1, Lpl*, and *Cst7* in mouse and *APOE*, *SPP1*, *CD81*, and *APOC1* in human, which are called disease-associated microglia (DAMs) or Alzheimer’s Disease (AD) microglia, although similar states have also been observed during normal aging^5,16,23^. We observed a population of DAM-like microglia in all patients, likely a result of advanced age and a history of neuropathology in some patients (Table S1). Indeed, the fraction of microglia expressing a DAM phenotype was highest in a patient in the cohort without COVID-19 who had an antemortem diagnosis of dementia. The relative abundance of this microglial state was indistinguishable between groups, arguing against the hypothesis that COVID-19 acutely drives the emergence of a DAM-like phenotype (Fig 1b).

To determine whether microglia from patients with COVID-19 exhibited a transcriptional signature distinct from patients who died from other causes, we performed pseudobulk differential expression analysis (DEA) on each major cell type cluster. This analysis revealed a pattern of gene expression in microglia from patients with COVID-19 that included a nearly complete downregulation of genes associated with cellular proliferation (*MKI67*, *CENPF*) and upregulation of the cell-cycle-arrest marker *CDKN1A* (encoding p21). While this pattern of gene expression has been referred to as immunosenescence, we did not detect significant upregulation of other senescence-associated genes (e.g., *CDKN2A*/p16 or *SERPINE1*/PAI-1; Fig. 1d-e).

Cytokine exposure has been suggested to drive the long-lasting cognitive impairment resulting from severe pneumonia caused by other pathogens that lack neurotropism, including influenza A viruses and bacteria^1–3,10,24,25^. Some of these cytokines, including TNF-α, IL-6, and IL-1β, can directly cross the blood-brain barrier and act on resident immune cells in the CNS, including microglia^26–28^. We examined whether microglia isolated from patients who died after SARS-CoV-2 infection exhibited an elevated response to any of these cytokines by comparing gene-set enrichment using MSigDB HALLMARK annotations^29^. Through group-wise gene-set enrichment analysis (GSEA) on pseudo-bulk data, we found that Hallmark TNF-α Signaling Via NF-κB (M5890) was the most significantly enriched gene set among all hallmark gene sets in MSigDB (q = 6.0×10^-16^; Fig. 1f). We further found through patient-wise gene module analysis that Hallmark TNF-α Signaling Via NF-κB (M5890) gene expression was significantly elevated in individuals who died after SARS-CoV-2 infection, suggesting that prolonged exposure to TNF-α or other NF-κB-activating cytokines may drive cell-cycle arrest in these patients (q = 3.6×10^-2^, Mann-Whitney; Fig. 1g). We confirmed the co-expression of three key marker genes of this cell state, *CDKN1A*, *CCL2*, and *IL1B*, in IBA1^+^ microglia with smFISH (RNAScope*™*) analysis of brain sections from the same patients and consistently observed cells co-expressing these markers in samples from patients who died with COVID-19 that were absent in control samples (Fig. 1h).

## COVID-19 is associated with greater cumulative systemic exposure to inflammatory cytokines compared to other causes of pneumonia

To determine if the transcriptomic changes in microglia we observed in patients who died after SARS-CoV-2 infection could have resulted from exposure to unusually high levels of inflammatory cytokines, we performed multiplexed profiling of 72 cytokines, 55 of which were of sufficient quality for downstream analysis (Supp. Fig. S3a-b). We analyzed serial plasma and alveolar samples collected by bronchoalveolar lavage (BAL) from 354 patients. These samples included patients with respiratory failure requiring mechanical ventilation for SARS-CoV-2 pneumonia (n = 93), pneumonia secondary to bacterial or fungal pathogens (n = 162), pneumonia resulting from other respiratory viruses (n = 41), conditions requiring mechanical ventilation for reasons unrelated to pneumonia (n = 45), and healthy controls (n = 13). Samples collected from all patients with respiratory failure were collected as part of an observational cohort study; samples from healthy controls were collected before the COVID-19 pandemic.

Comprehensive demographic data from these cohorts are available in Supp. Table S2. Findings from this cohort have been previously reported^7,30,31^. Patients with severe SARS-CoV-2 pneumonia were similar to other groups of mechanically ventilated patients in age, sex, severity of illness as measured by the mean daily Sequential Organ Failure Assessment (SOFA; q ≥ 0.05, Mann-Whitney), Acute Physiology Score (APS; q ≥ 0.05, Mann-Whitney), and mortality (q ≥ 0.05; chi–square test of proportions); however, patients with non-viral pneumonias (“Other Pneumonia”) were older (q = 2.6×10^-2^, Mann-Whitney) and had higher SOFA scores (q = 1.3×10^-3^, Mann-Whitney) and APS scores (q = 4.3×10^-2^, Mann-Whitney). Patients with SARS-CoV-2 pneumonia were more likely to self-describe as Hispanic or Latino than all other groups (q < 0.05, chi-square test of proportions) and had higher BMIs (q < 0.05, Mann-Whitney) compared with the rest of the cohort. Despite similar severity of illness on presentation and hospital mortality rate, the duration of mechanical ventilation and ICU stay was 1.8–2.4-fold longer in patients with SARS-CoV-2 pneumonia compared with all other groups of mechanically ventilated patients (q < 0.05, Mann-Whitney; Supp. Data File 2). In accordance with our previous findings, analysis of BAL fluid samples collected from all patients other than healthy controls revealed an elevated percentage of lymphocytes from patients with COVID-19 relative to all other groups of mechanically ventilated patients (q < 0.05, Mann-Whitney; Suppl. Fig. S2). One hundred eighty-seven patients in the cohort had BAL samples collected within 48 hours of intubation (early). Severity of illness scores and mortality rates were similar in patients with early BAL samples compared with the entire cohort of mechanically ventilated patients (q ≥ 0.05, Mann-Whitney and q ≥ 0.05, chi-square test of proportions, respectively; Supp. Data File 3).

We proposed a model in which the relatively long clinical course of patients with COVID-19 results from spatially-restricted inflammatory circuits between alveolar macrophages harboring SARS-CoV-2 and activated T cells in the alveolar space^7,32^. This model has since been confirmed by other groups^33–36^. Enhanced transcription of chemokines promoting chemotaxis of monocytes and T cells, including *CXCL10*, *CCL8*, and *CCL2* by monocyte-derived alveolar macrophages infected with or harboring SARS-CoV-2 is key to this model. In support of this model, samples collected within 48 hours of intubation from patients with COVID-19 clustered distinctly from other patient groups, driven by CXCL10 and CCL8 in BAL samples and CXCL10 in plasma (Fig. 2a-b). Elevated concentrations of CCL2 and CCL8 were also observed in BAL samples from patients with COVID-19, relative to patients with pneumonia secondary to nonviral pathogens (q < 0.05, Mann-Whitney; Fig. 2c-d). Indeed, concentrations of CXCL10 were significantly higher in the first 48 hours of intubation in both BAL fluid and plasma from patients with COVID-19, relative to all other groups (q < 0.05, Mann-Whitney). Levels of CCL2 and CCL8 in BAL fluid as well as CXCL10 in plasma samples collected during the first 48 hours of intubation were higher in patients with COVID-19 relative to all other groups, with the exception of patients with other viral pneumonias (q < 0.05, Mann-Whitney; Fig. 2c-d).

**Figure 2.**
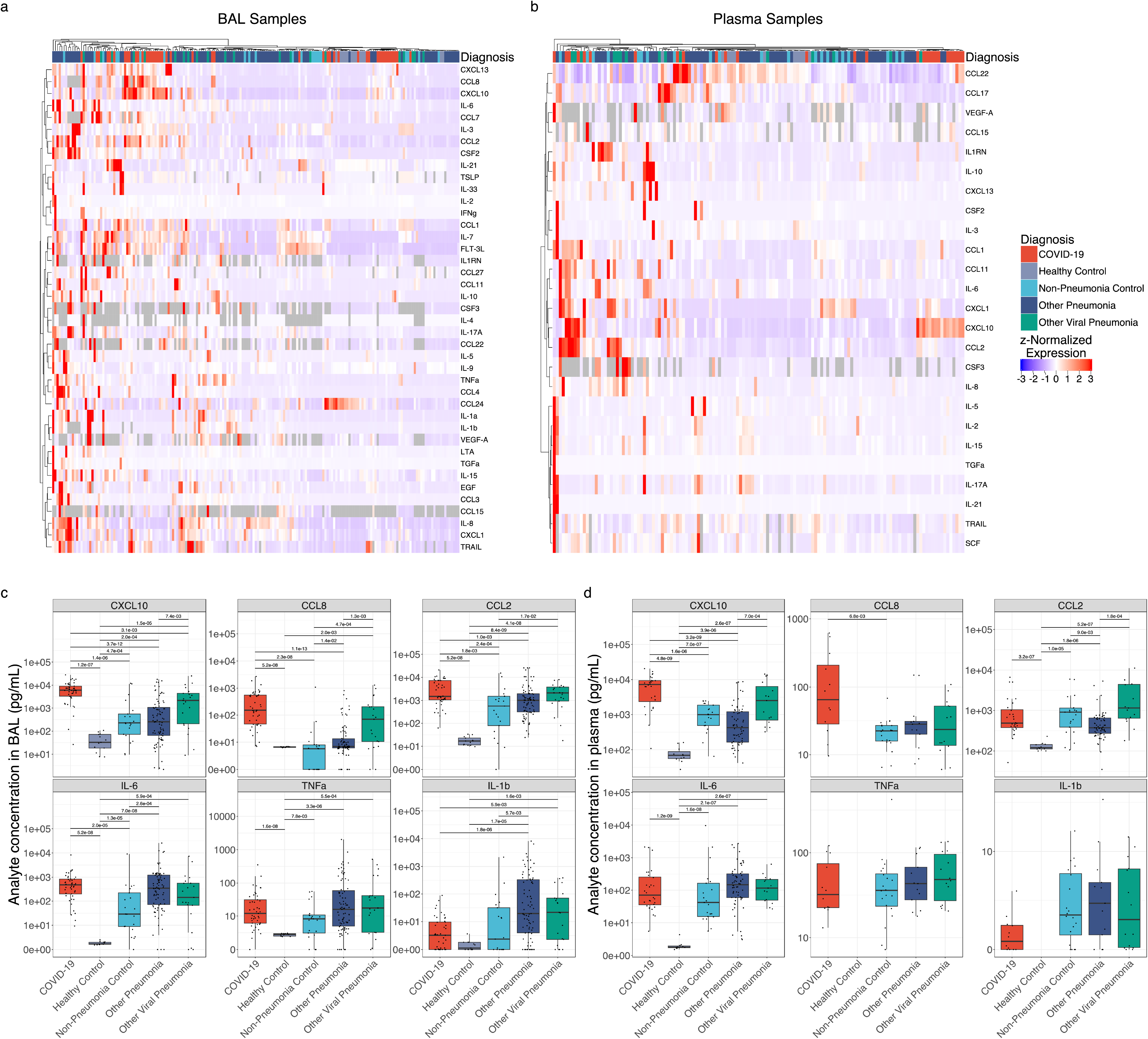
COVID-19 is distinguished from pneumonias of similar severity by expression of T cell chemokines. **(a)** Hierarchical clustering of 41 cytokines showing significant variability by diagnosis (q < 0.05, Kruskal–Wallis) from 187 BAL samples collected in the first 48 hours of intubation from 183 patients with an early BAL. **(b)** Hierarchical clustering of 25 cytokines showing significant variability by diagnosis (q < 0.05, Kruskal–Wallis) from 137 early plasma samples from 134 patients. **(c)** Expression of COVID-19-defining T lymphocyte and monocyte chemokines and key pro-inflammatory cytokines from 479 BAL samples collected throughout the duration of mechanical ventilation from 332 patients. **(d)** Expression of COVID-19-defining T lymphocyte and monocyte chemokines and key pro-inflammatory cytokines from 396 plasma samples collected throughout the duration of mechanical ventilation from 262 patients.

Concentrations of other BAL fluid cytokines in samples collected during the first 48 hours of intubation, including CXCL1, IFNɣ, IL-1β, IL-6, IL-8, and TNF-ɑ, were higher in patients with pneumonia relative to healthy controls, but were largely similar between groups of mechanically ventilated patients (complete comparisons are included in Extended Data 1). A similar pattern was observed in the plasma of patients with COVID-19 compared to patients with other causes of pneumonia and respiratory failure, arguing against an unusually severe inflammatory response or “cytokine storm” in patients with COVID-19 (Fig. 2d)^37,38^. Consistent with previous findings, the concentrations of IL-1β, IL-6, and TNF-ɑ were higher in all groups of mechanically ventilated patients compared with healthy controls (Fig. 2c-d)^37^.

We wondered whether the roughly 2-fold increase in the duration of illness in patients with COVID-19 might result in a higher cumulative exposure to pro-inflammatory cytokines. As the peak levels of inflammatory cytokines did not differ between patients with COVID-19 compared with similarly ill patients with pneumonia secondary to other pathogens, cumulative exposure could only be higher if the levels of inflammatory cytokines did not normalize over the course of the illness. We therefore performed geometric integration of cytokine expression over the ICU course, yielding a single value corresponding to an estimate of cumulative exposure to a given cytokine during the ICU stay (Fig. 3e). Strikingly, samples originating from patients with COVID-19 – particularly patients with unfavorable outcomes – clustered together and were defined by greater cumulative exposure to pro-inflammatory cytokines both in BAL fluid and plasma (Fig. 3a-b). Among the many cytokines with significantly greater cumulative exposure in COVID-19 were CXCL10, CCL8, CCL2, IL-6, and TNF-α (Fig. 3c-d; q < 0.05, Mann-Whitney).

**Figure 3.**
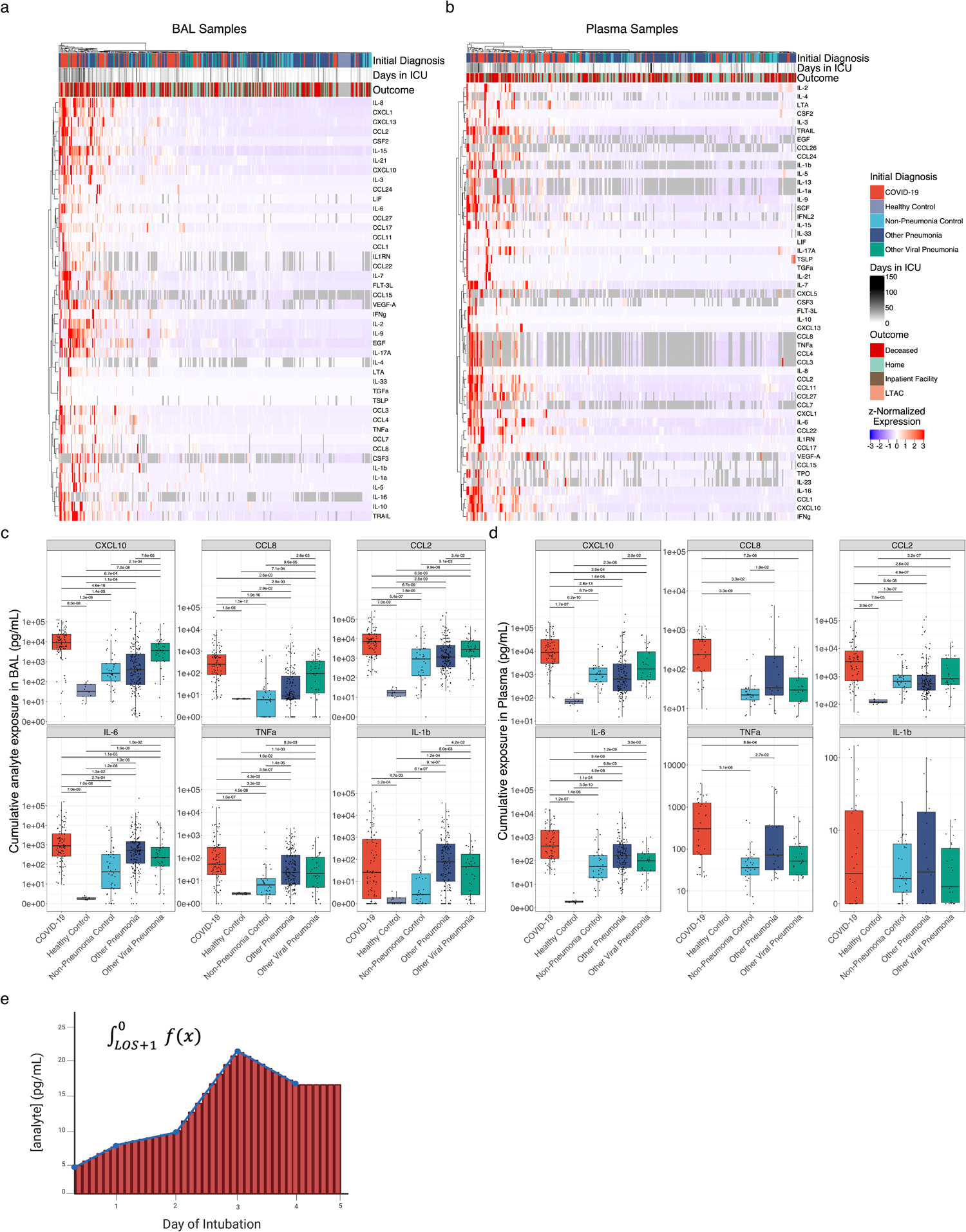
Cumulative but not peak exposure to pro-inflammatory cytokines is higher in patients with severe SARS-CoV-2 pneumonia compared to patients with severe pneumonia secondary to other pathogens. **(a)** Hierarchical clustering of cumulative exposure to 44 BAL cytokines showing significant variability by diagnosis (q < 0.05, Kruskal–Wallis) from 327 patients estimated by geometric integration of the levels of 479 BAL samples collected throughout the duration of mechanical ventilation. **(b)** Hierarchical clustering of cumulative exposure to 51 plasma cytokines showing significant variability by diagnosis (q < 0.05, Kruskal–Wallis) from 258 patients estimated by geometric integration of the levels of 396 plasma samples collected throughout the duration of mechanical ventilation. **(c)** Cumulative expression of selected pro-inflammatory cytokines in BAL fluid from (a). **(d)** Cumulative expression of selected pro-inflammatory cytokines in plasma from (b). **(e)** Schematic for calculation of cumulative exposure for each cytokine assayed for each patient throughout the course of mechanical ventilation by geometric integration. BAL samples from 3 patients and plasma samples from 2 patients receiving long-term mechanical ventilation were excluded from these analyses.

## Levels of T cell and monocyte chemoattractants are lower in patients with severe COVID-19 who received corticosteroid treatment

Prior to publication of the RECOVERY Collaborative study demonstrating efficacy of dexamethasone treatment in patients with severe COVID-19^6^, there was clinical equipoise around the prescription of systemic corticosteroids as a therapy for these patients. We took advantage of the inconsistent use of corticosteroids before this trial to examine their effect on cytokine expression in BAL fluid and plasma. We performed unbiased comparison of the concentrations of all 55 analytes as a function of steroid treatment. In BAL fluid, only the levels of CCL8, CXCL10, and CCL7 were significantly lower, and the levels of IL-10 were higher, in patients with COVID-19 after they received corticosteroids compared to those who did not receive corticosteroids (q < 0.05, Mann-Whitney). In plasma, only the concentrations of CXCL10 and CCL7 were significantly lower, and IL-10 was significantly higher, after patients received corticosteroids (q < 0.05, Mann-Whitney). Notably, while these cytokines were detected in all groups of intubated patients, differences in the concentration of these cytokines as a function of corticosteroid treatment were only observed in patients with COVID-19 (q ≥ 0.05, Mann-Whitney; Fig. 4a). In previously published scRNA-seq data of BAL fluid from this cohort, the expression of *CXCL10*, *CCL2*, *CCL7*, and *CCL8* was highest in monocyte-derived alveolar macrophages harboring SARS-CoV-2^7^.

**Figure 4.**
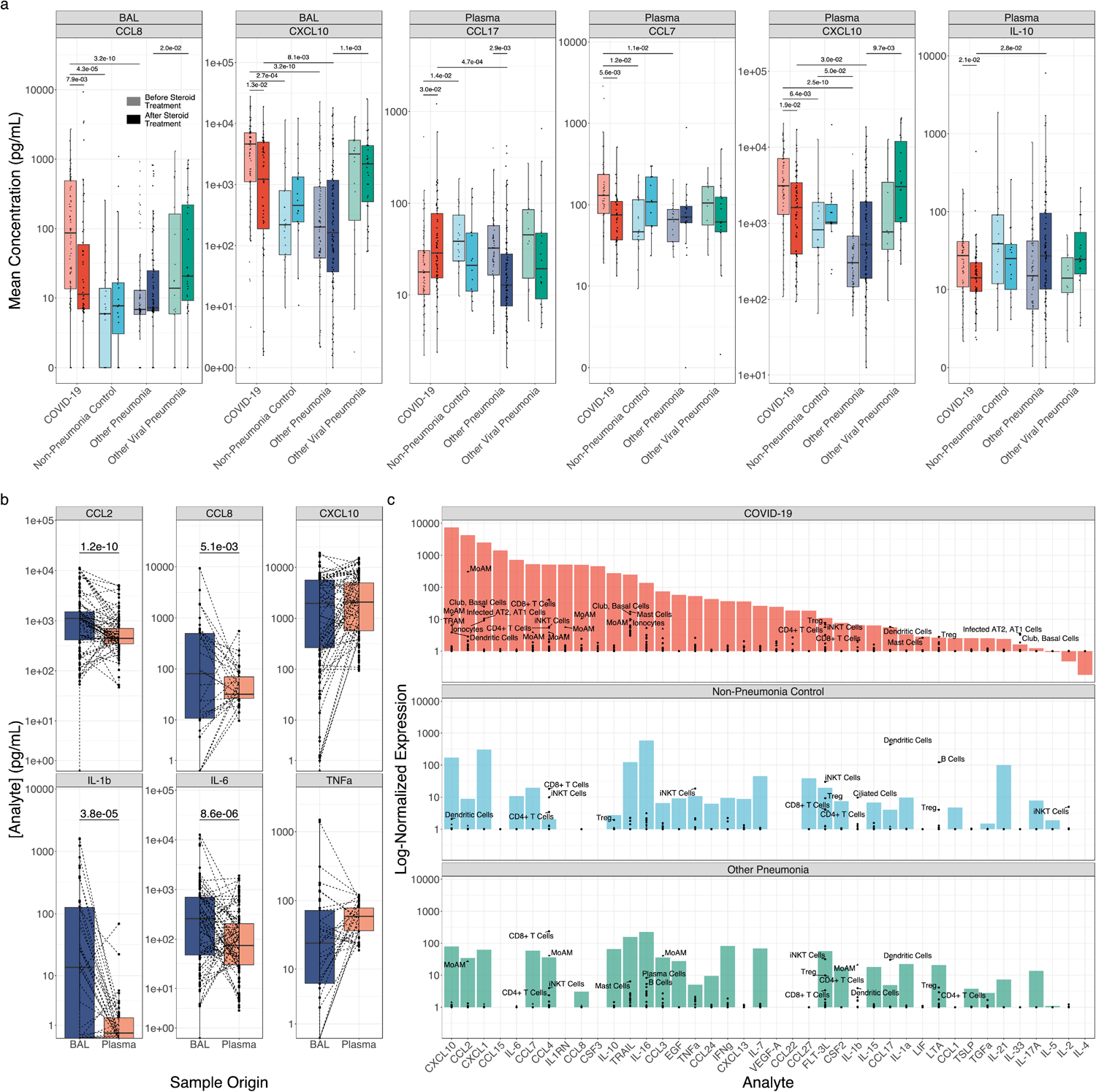
Corticosteroid treatment is associated with reductions in T cell chemokine expression in monocyte-derived alveolar macrophages. **(a)** Boxplots of cytokine expression for all BAL fluid and plasma cytokines exhibiting significantly altered expression (q < 0.05, Mann-Whitney) following corticosteroid treatment. Lightly shaded boxes represent cytokine expression values prior to corticosteroid treatment, and darkly shaded boxes represent expression values after corticosteroid treatment. **(b)** Paired comparisons of cytokine expression in BAL and plasma for all paired samples (paired Mann-Whitney). **(c)** Deconvolution of “bulk” cytokine expression in BAL fluid by scRNA-seq of cells isolated from BAL fluid. Mean cytokine gene expression for each cell type detected in scRNA-seq data from^7^ (black points) is overlaid on bulk cytokine expression by multiplexed cytokine array (filled bars) to identify cell type contributors to cytokine expression.

## Circulating cytokines in patients with COVID-19 originate from the alveolar space

We reasoned that if plasma cytokines in patients with SARS-CoV-2 originated from the alveolar space, we should observe a correlation between the concentration of BAL fluid cytokines and the concentration of plasma cytokines. When comparing all analytes for all paired BAL and plasma samples, we observed a significant, nonlinear correlation (ρ = 0.49, P < 2.2×10^-16^, Spearman rank correlation). Correlations were particularly strong for key cytokine markers of SARS-CoV-2 infection, including CXCL10 (ρ = 0.55, q = 2.1×10^-25^, Spearman rank correlation), CCL2 (ρ = 0.52, q = 3.1×10^-22^, Spearman rank correlation), CCL8 (ρ = 0.41, q = 1.6×10^-5^, Spearman rank correlation), and IL-6 (ρ = 0.38, q = 5.3×10^-11^, Spearman rank correlation). We also compared the concentrations of inflammatory cytokines in paired BAL fluid and plasma samples. Despite the 10-100 fold dilution of alveolar fluid by the BAL procedure^39^, measured concentrations of many inflammatory cytokines in BAL fluid, including IL-6, IL-1β, CCL2, and CCL8, exceeded the concentrations in plasma, while the levels of CXCL10 and TNF-α were similar (Fig. 4b). These data suggest the alveolus is a major contributor to pro-inflammatory cytokine levels in plasma during COVID-19 and are consistent with the known tropism of SARS-CoV-2 for the respiratory epithelium and alveolar macrophages^40^.

We then used previously published scRNA-seq data from BAL fluid obtained from 10 patients in this cohort to identify candidate cells in the lung that might be the source of inflammatory cytokines^7^. Monocyte-derived alveolar macrophages expressed high levels of *CXCL10*, *CCL8*, *CCL2*, *CCL3*, and *IL1RN*. Surprisingly, while IL-6 concentrations were higher in BAL fluid from patients with COVID-19, and these concentrations were strongly correlated with plasma concentrations, we did not identify cells expressing *IL6* in BAL fluid, suggesting this cytokine is produced by cells in the lung parenchyma that are not sampled by the BAL procedure (Fig. 4c).

## Discussion

Neurologic and psychiatric symptoms are among the most common complaints in patients with PASC^4,9–11^. Even more concerning, survivors of severe pneumonia, particularly the elderly, are at an increased risk of dementia for years after hospitalization^1–3^. While some studies have suggested these symptoms result from direct infection of the CNS by SARS-CoV-2, rigorous studies of clinical samples and autopsy tissues from patients who died from COVID-19 reveal that SARS-CoV-2 infection is limited to the airway epithelium and alveolar macrophages^7,11,19,20,41,42^. In instances when SARS-CoV-2 virus was detected in the brain, it was limited to the hypothalamus, which is anatomically adjacent to the nasopharynx and has a relatively permeable blood brain barrier, raising the question of artifactual contamination during tissue processing.^43^ Indeed, careful studies of the mechanisms underlying the loss of taste and smell in patients with COVID-19 failed to find evidence of infection in olfactory neurons, instead implicating inflammatory signals from adjacent infected nasopharyngeal epithelia^44,45^. In flow cytometry-sorted immune cells from the cortex of a small cohort of patients who died after a diagnosis of COVID-19, we did not detect direct CNS infection with SARS-CoV-2 or reactivation of EBV1. Instead, we saw a transcriptional phenotype in microglia that included downregulation of genes associated with proliferation, upregulation of *CDKN1A*, and higher expression of inflammatory genes associated with signaling through TNF-α. Consistent with this finding, cumulative exposure to TNF-α and other cytokines originating from the lungs of patients with COVID-19 was higher relative to patients with pneumonia from other pathogens. As microglial inflammation has been demonstrated to reduce neuronal plasticity, synapse density, and memory formation^46–50^, these changes may partly explain deficits in executive function observed in COVID-19 survivors^9,13^.

Whether the cognitive changes and increased rates of dementia observed in patients with COVID-19 are more severe or frequent than those observed in survivors of pneumonia secondary to other pathogens is unknown^1–3^. Consistent with another report, we found the levels of inflammatory cytokines originating in the lung at the time of peak illness severity (within 48 hours of intubation for respiratory failure) were similar in patients with COVID-19 compared to patients with pneumonia secondary to other pathogens^51^. We took advantage of serial sampling in our cohort to show that the levels of lung-derived inflammatory cytokines remain elevated in patients with COVID-19 over the course of their ICU stay. As we and others have reported, the duration of critical illness is twice as long in patients with COVID-19 compared to patients with pneumonia secondary to other pathogens^7,32,52–56^. The data reported here show that patients with COVID-19 have a higher cumulative systemic exposure to lung-derived inflammatory cytokines during their illness and suggest a “cytokine monsoon” rather than “cytokine storm” might drive more severe or prolonged post-acute sequelae of infection in COVID-19 survivors through prolonged activation of NF-κB.

Using data obtained from the analysis of alveolar samples, we developed a model that explains the long duration of illness in patients with COVID-19, which has been confirmed by other groups^33–36^. Key to this model is the presence of self-sustaining inflammatory circuits between alveolar macrophages harboring or infected with SARS-CoV-2 and activated T cells in the alveolar space that are maintained by the release CXCL10, CCL8, and CCL2. Our findings support this model by showing alveolar and plasma levels of these cytokines differentiate patients with COVID-19 from patients infected with other pathogens. Moreover, we found that corticosteroids, which are effective in SARS-CoV-2 but are not universally effective in all causes of pneumonia, were associated with lower levels of these inflammatory cytokines in the lung and plasma, possibly by targeting their expression from alveolar macrophages harboring SARS-CoV-2^6,57^.

Our study has limitations. Most importantly, while our sampling of alveolar fluid and plasma includes the largest cohort of patients reported to date, our analysis of cortical tissue includes only a small number of patients and a single anatomical region. It is therefore possible that microglial phenotypes that develop in a minority of patients were missed in our analysis or that our small cohort represents outliers. Additionally, other brain regions may demonstrate distinct patterns of resident immune cell activation. Second, while our data suggest the transcriptomic changes in microglia we observe in patients with COVID-19 are induced by TNF-α and other cytokines, we cannot infer causality from these observational data. Third, the administration of corticosteroids in our observational cohort was not randomized and did not include a placebo control. We therefore cannot conclude the differences in cytokine expression we observed are causally linked to steroid administration. Fourth, as our scRNA-seq data from BAL fluid does not effectively capture all lung cell types, we cannot determine if alveolar macrophages are the primary contributors to the expression of chemokines that attract and activate T cells and monocytes. Finally, several lines of evidence suggest SARS-CoV-2 induces nonproductive infection of nasopharyngeal and alveolar macrophages^40,58^. Nevertheless, human alveolar macrophages do not express ACE2 (the receptor for SARS-CoV-2) and we cannot definitively distinguish infection from uptake^59, 60^. Irrespective of whether alveolar macrophages in patients with COVID-19 are infected by or harbor SARS-CoV-2, our published data suggest they exhibit distinct transcriptional responses when compared to SARS-CoV-2-negative cells.

---

**Figure S1.**
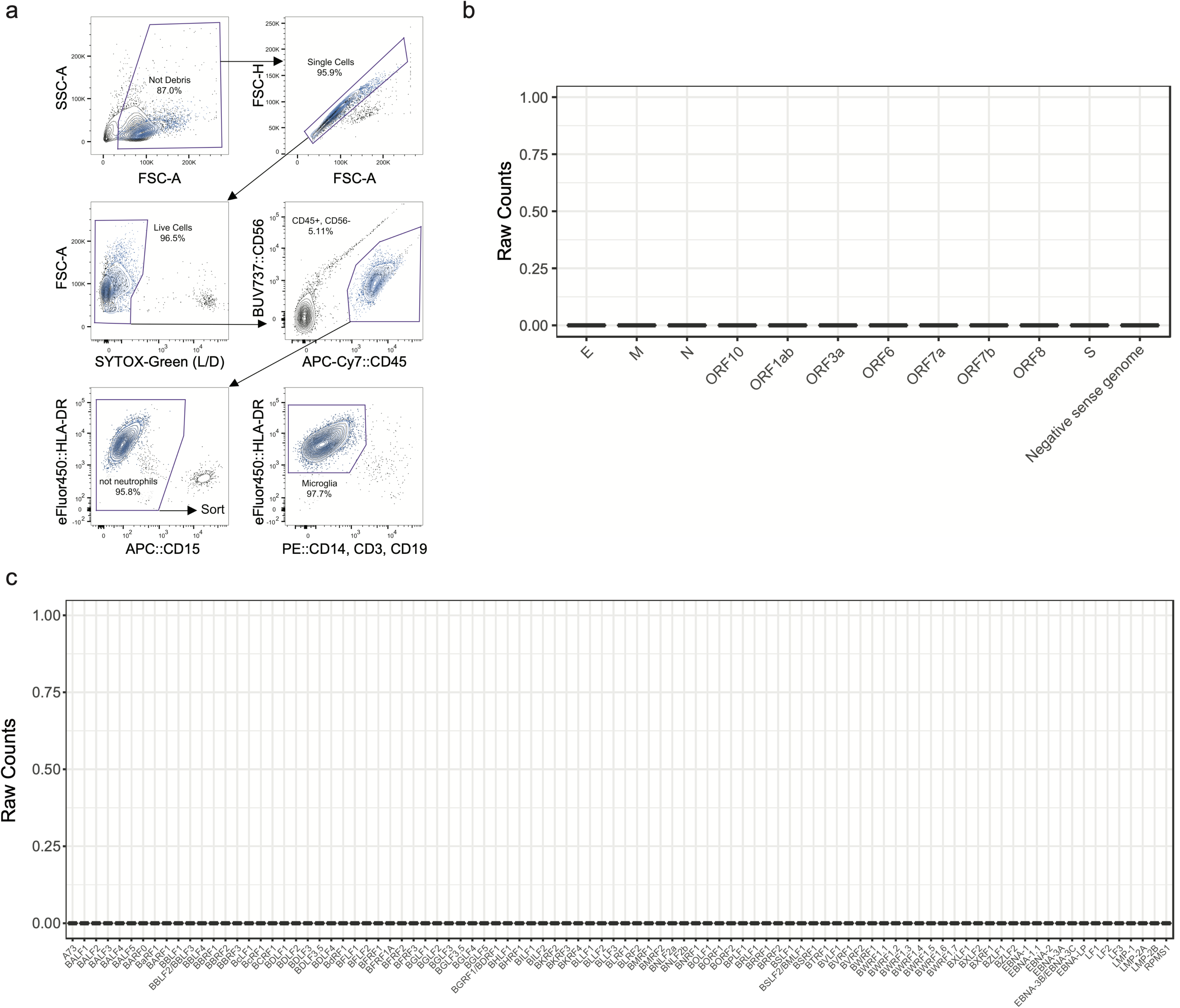
Related to figure 1. **(a)** Representative flow cytometry sorting scheme for isolating neuro-immune cells from human postmortem brain tissue from a frontal lobe sample. **(b)** Boxplots of raw gene counts for all genes in the SARS-CoV-2 (NC045512.2) genome from human neuro-immune cell scRNA-seq. Zero counts are detected for all genes. **(c)** Boxplots of raw gene counts for all genes in the EBV1 genome (NC007605.1) from human neuro-immune cell scRNA-seq. Zero counts are detected for all genes.

**Figure S2.**
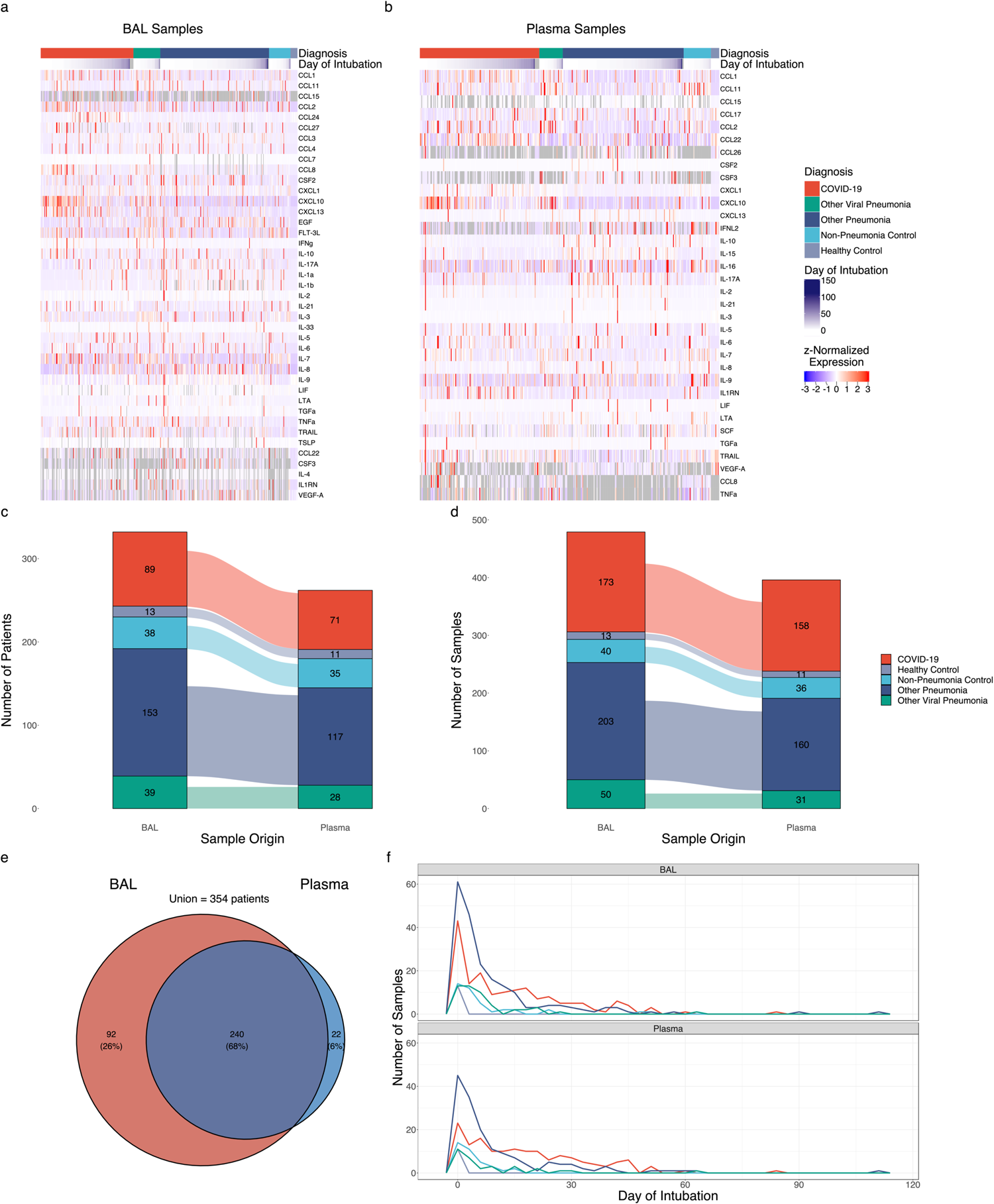
Related to figure 2. **(a)** Heatmap of 41 cytokines showing significant variability by diagnosis (q < 0.05, Kruskal–Wallis) from 479 BAL samples collected throughout the entire ICU stay from 332 patients. **(b)** Heatmap of 31 cytokines showing significant variability by diagnosis (q < 0.05, Kruskal–Wallis) from 396 plasma samples collected throughout the entire ICU stay from 262 patients. **(c,d)** Alluvial plot of total numbers of patients included in the BAL fluid (c) and plasma (d) datasets. **(e)** Venn diagram of patients with BAL and plasma samples. **(f)** Frequency polygon plot of BAL and plasma sampling distributions throughout patient stay by group.

**Figure S3.**
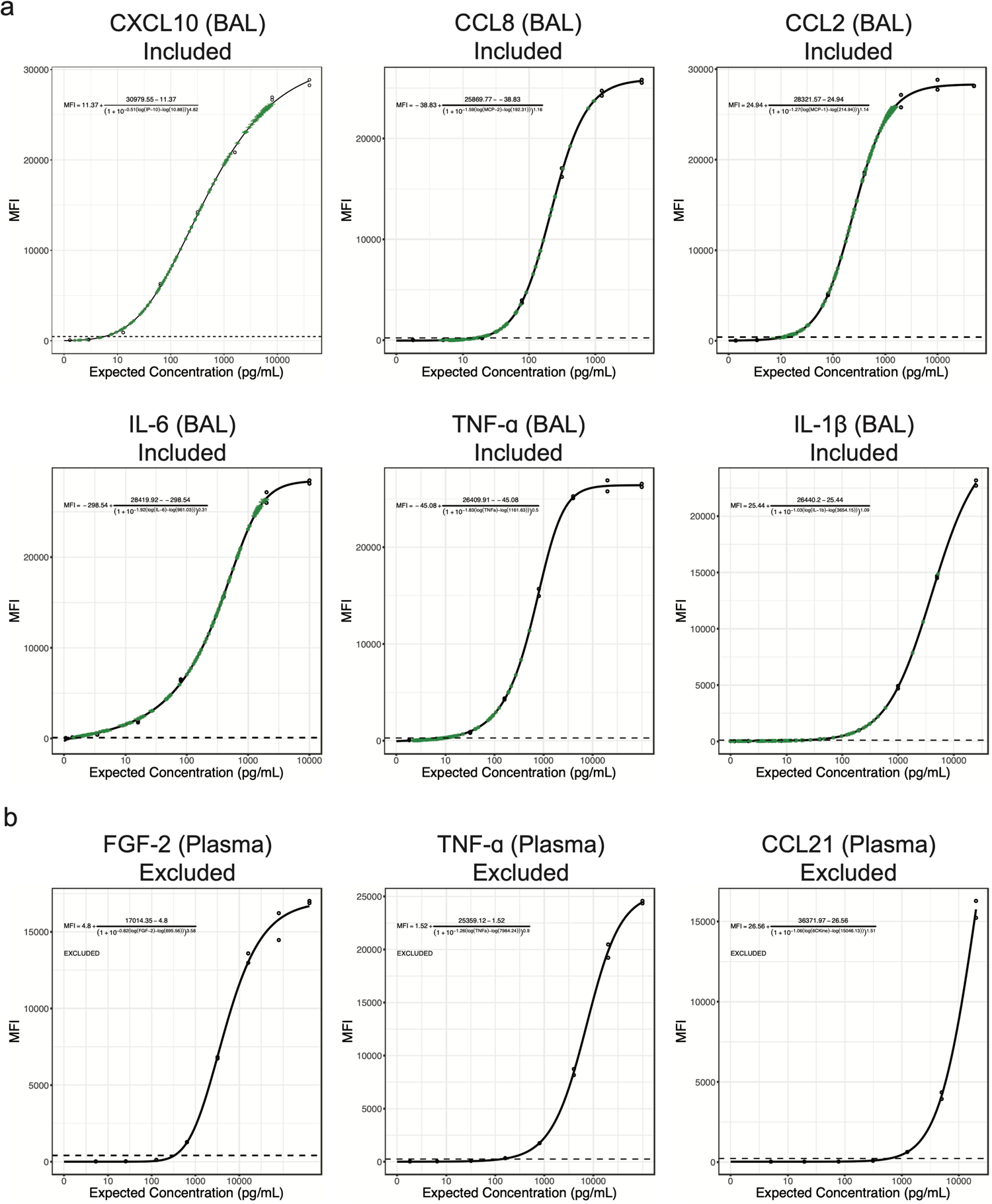
Representative standard curve fitting plots for cytokine data exclusion. (a) Representative standard curves and fitted cytokine expression values for analytes included in this study. From left to right, top to bottom: CXCL10 (IP-10), CCL8 (MCP-2), CCL2 (MCP-1) IL-6, TNF-ɑ, IL-1b. **(b)** Representative standard curves for analytes excluded for poor sensitivity in this study. From left to right: FGF-2, TNF-ɑ, and CCL21. For all plots, empty points represent standards of known concentration used for curve-fitting. Black curves and associated formulae represent the function fitted to these points. Green points represent experimental values, and green whiskers are the standard error of prediction. Dotted lines are the upper limit of the 95% confidence interval of lower asymptote prediction.

**Supplemental Data File 1.** Pseudobulk DESeq2 differential expression analysis results from human microglia scRNA-seq in COVID-19 vs. controls.

**Supplemental Data File 2.** Complete pairwise comparisons between diagnosis groups for all relevant cytokine cohort metadata (see Table 2).

**Supplemental Data File 3.** Complete comparisons between patients with early BALs and the entire cohort for all relevant cytokine cohort metadata.

**Supplemental Data File 4.** List of the complete NU SCRIPT investigators.

## Data availability

The complete scRNA-seq dataset, including raw FASTQ files, raw and normalized counts, and all relevant metadata will be made available through dbGAP by final publication. Cytokine expression and cumulative cytokine expression values including all relevant metadata will be made available through PhysioNet by final publication. An interactive version of Figure 1a with all gene expression data and relevant metadata is available at https://nupulmonary.org/covid-19/human_microglia/?ds=human_microglia_COVID-19.

## Code availability

The complete code used to process data and generate all figures is available at https://github.com/NUPulmonary/2023_Grant_Poor. Scripts used for data processing are available at https://github.com/NUPulmonary/utils.

## Acknowledgments

We would like to thank Constadina Arvanitis in the Northwestern University Center for Advanced Microscopy & Nikon Imaging Center for substantial help with optimization of combined RNA-scope and immunofluorescence on human brain samples. Imaging work was performed at the Northwestern University Center for Advanced Microscopy generously supported by CCSG P30 CA060553 awarded to the Robert H Lurie Comprehensive Cancer Center. We further thank Cara J. Gottardi for her insightful comments on smFISH image presentation and analysis. Finally, we thank the many human subjects and their families for their invaluable contributions to this research. The Genomics Compute Cluster is part of Quest, Northwestern University’s high-performance computing facility, with the purpose to advance research in genomics. Single-cell RNA-seq was performed with support from Simpsons Querrey Institute for Epigenetics. Northwestern University Flow Cytometry Core Facility and the Northwestern University Metabolomics Core Facility are supported by NCI Cancer Center Support Grant P30 CA060553 awarded to the Robert H. Lurie Comprehensive Cancer Center. Cell sorting was performed on a BD FACSAria SORP cell sorter purchased through the support of NIH 1S10OD011996-01.

The authors acknowledge the support of The Simpson Querrey Lung Institute for Translational Sciences (SQLIFTS) at Northwestern University and the support of the Dixon Translational Research Grants Initiative at Northwestern Medicine and the Northwestern University Clinical and Translational Sciences Institute (UL1TR001422). This work was also supported by the Chicago Biomedical Consortium with Support from the Searle Funds at The Chicago Community Trust. R.A.G was funded by NIH grants T32AG020506 and 1F31AG071225. T.A.P. was supported by NIH grant T32HL076139. J.I.B. was supported by 5T32HL076139 and UL1TR001422. R.M.T. was supported by NIH grants R01ES034350 and R01ES027574. R.G.W. is supported by NIH grants U19AI135964, U01TR003528, P01HL154998, R01HL14988, and R01LM013337. B.D.S was supported by NIH grants R01HL149883, R01HL153122, P01HL154998, P01AG049665, and U19AI135964. A.V.M was supported by NIH grants U19AI135964, P01AG049665, P01HL154998, R01HL153312, R01HL158139, R01ES034350, R21AG075423. G.R.S.B was supported by a Chicago Biomedical Consortium grant, Northwestern University Dixon Translational Science Award, Simpson Querrey Lung Institute for Translational Science (SQLIFTS), and NIH grants AG049665, HL154998, HL14575, HL158139, HL147290, AG075423, AI135964 and The Veterans Administration award I01CX001777.

## Materials and Methods

### Human subjects (BAL and plasma collection)

All human subjects research was approved by the Northwestern University Institutional Review Board. Samples from patients with COVID-19, viral pneumonia, other pneumonia and non-pneumonia controls were collected from participants enrolled in Successful Clinical Response In Pneumonia Therapy (SCRIPT) study STU00204868. Data from this cohort has been published previously and is available from dbGAP (phs002300.v1.p1)^7,30,31^. All subjects or their surrogates provided informed consent. Patients ≥ 18 years of age with suspicion of pneumonia based on clinical criteria (including but not limited to fever, radiographic infiltrate, and respiratory secretions) were screened for enrollment into the SCRIPT study. Inability to safely obtain BAL or NBBAL were considered exclusion criteria ^61^. In our center, patients with respiratory failure are intubated based on the judgment of bedside clinicians for worsening hypoxemia, hypercapnia, or work of breathing refractory to high-flow oxygen or non-invasive ventilation modes. All patients were admitted to Northwestern Memorial Hospital in Chicago between June 15, 2018 and September 29, 2021. Bronchoscopy was most commonly performed as part of routine clinical care to guide antimicrobial therapy, with paired blood draws for plasma for most samples. Extubation occurs based on the judgment of bedside clinicians following a trial of spontaneous breathing in patients demonstrating physiologic improvement in their cardiorespiratory status during their period of mechanical ventilation. Management of patients with COVID-19 was guided by protocols published and updated on the Northwestern Medicine website as new information became available over the pandemic. Clinical laboratory testing including studies ordered on bronchoalveolar lavage fluid was at the discretion of the care team, however, quantitative cultures, multiplex PCR (BioFire Film Array Respiratory 2 panel), and automated cell count and differential were recommended by local ICU protocols. Most patients also underwent urinary antigen testing for Streptococcus pneumoniae and Legionella pneumophila on admission. Clinicians were encouraged to manage all patients, including those with COVID-19, according to ARDSnet guidelines including the use of a higher PEEP/lower FIO2 strategy for those with severe hypoxemia^62,63^. Prone positioning (16 hours per day) was performed in all patients with a PaO2/FiO2 <150 who did not have contraindications^64^. In those who had a response to prone positioning evident by improved oxygenation, prone positioning was repeated. Esophageal balloon catheters (Cooper Surgical) were placed at the discretion of the care team to estimate transpulmonary pressure and optimize PEEP, particularly in patients with a higher than normal BMI. Pneumonia category adjudication was performed by five critical care physicians using a published protocol^65^. Clinical laboratory data were obtained from the Northwestern Medicine Enterprise Data Warehouse using Structured Query Language (SQL). APS and SOFA scores were generated from the Electronic Health Record using previously validated programming. Anonymized clinical data from this cohort is available on Physionet (https://doi.org/10.13026/5phr-4r89)^30,31^. A complete list of the investigators involved in this study is available in Supp. Data File 4.

### NBBAL and BAL procedures

Consent was obtained from patients or legal decision makers for the bronchoscopic procedures. Bronchoscopic BAL was performed in intubated ICU patients with flexible, single-use Ambu aScope (Ambu) devices. Patients were given sedation and topical anesthetic at the physician proceduralist’s discretion. Vital signs were monitored continuously throughout the procedure. The bronchoscope was wedged in the segment of interest based on available chest imaging or intraprocedure observations, aliquots of 30 cc of normal saline at a time, generally 90–120 cc total, were instilled and aspirated back. The fluid returned following the first aliquot was routinely discarded. Samples were split (if sufficient return volume was available) and sent for clinical studies and an aliquot reserved for research. A similar procedure was applied to non-bronchoscopic BAL (NBBAL); however, NBBAL was performed with directional but not visual guidance, and as usual procedural care by a respiratory therapist rather than a pulmonologist^61^. For the bronchoscopies performed in the COVID-19 patients, additional precautions were taken to minimize the risk to healthcare workers including only having essential providers present in the room, clamping of the endotracheal tube, transient disconnection of the inspiratory limb from the ventilator, and preloading of the bronchoscope through the adapter^64^; sedation and neuromuscular blockade to prevent cough was administered for these procedures at the physician’s discretion^66^. In most cases the early bronchoscopy was performed immediately after intubation^61^.

### BAL procedures (Duke University healthy controls)

Healthy volunteers were enrolled in the study Pro00088966 and Pro00100375 at Duke University. Bronchoscopic BAL was performed in patients in the bronchoscopy suite or in the intensive care unit. Patients were given sedation and topical anesthesia at the discretion of the physician performing the bronchoscopy. The most involved bronchopulmonary segment was identified based on clinician based on review of the chest CT scan and 90–120 ml of saline was instilled into the segment of interest and aspirated back with the first 5 cc of return discarded.

### Plasma collection

Patient whole blood was collected in lithium heparin tubes on the same day BAL or NBBAL procedures were performed. The cellular fraction was spun down for 10 minutes at 1690 x g at 4°C, and the plasma fraction was removed and stored at −80°C prior to multiplexed cytokine analysis.

### Human brain autopsy

Autopsy was performed at Northwestern Memorial Hospital, as approved under IRB STU212579 and CSRC-1661. Postmortem interval for all samples is reported in Suppl. Table S1. During routine brain autopsy, sections of the frontal cortex were removed by dissection and placed in sufficient pre-cooled HypoThermosol solution (BioLife Solutions 101104) to cover. Under aseptic conditions, any remaining arachnoid mater was removed on ice and discarded. Samples were then divided into 2 sections. The minor section was fixed in ice-cold fresh 4% formaldehyde (Electron Microscopy Sciences 15714) in 1X PBS for 48-72 hours, before being transferred to 1X PBS (Corning 21-040-CM) + 0.01% sodium azide (Sigma S2002) indefinitely. The major section was chopped into ∼3mm strips, stored at 4°C in HypoThermosol FRS (BioLife Solutions 101102), and processed for flow cytometry sorting as described below 0-48 hours later.

### Human brain tissue processing and isolation of single-cell suspension

Free-floating sections were placed on ice and rinsed briefly with 1X HBSS (Fisher Scientific 21023CM) and strained. Tissues were then chopped thoroughly in 1mL ice-cold digest buffer consisting of 1X Papain Dissociation System (Fisher NC9212788; 1 vial dissolved in 5mL HBSS to yield 20U of papain/mL in 1mM L-cysteine with 0.5mM EDTA) and 1mg/mL DNAse I (Roche 10104159001) with curved scissors. Chopped tissue was then transferred to gentleMACS C-tubes (Miltenyi 130-093-237) and mixed with 1mL HBSS. Samples were then mechanically dissociated using a gentleMACS Octo Dissociator using the stock program “m_brain_03_01”. Samples were then shaken at 200rpm, 37°C for 30 minutes, followed by a second round of mechanical dissociation. Digestion was then stopped by mixing samples with 18mL ice-cold, sterile-filtered 1% BSA (Sigma SLBW2268) in 1X HBSS. Cell suspensions were then mashed through a 70-µm filter with 3×10mL ice-cold 1% BSA in HBSS into a fresh 50mL conical tube. The resulting single-cell suspension was then pelleted at 400*g* for 10 minutes at 4°C and resuspended in 25mL RT 30% Percoll (Millipore-Sigma GE17-0891-01) in 1X HBSS without calcium or magnesium (Gibco 14185-052). The resultant suspension was then slowly layered on top of 5mL 70% Percoll in 1X HBSS without calcium or magnesium. Density centrifugation was performed at 600*g*, brake: 0, acceleration: 4 for 30 minutes at room temperature. Myelin and debris were removed using a vacuum apparatus, and 5-10mL of the cell-containing interphase was transferred to a fresh 50-mL conical tube, discarding the RBC/debris pellet. The purified cell suspension was then diluted 3:1 in ice-cold 1X HBSS and pelleted at 400*g* for 5 minutes at 4°C. Cells were then resuspended in 15mL ice-cold 1x HBSS and again pelleted at 400*g* for 5 minutes at 4°C and resuspended in 500µL ice-cold MACS buffer (Miltenyi 130-091-221). A 10-µL aliquot was then mixed with 10µL 2X AOPI (Nexcelom NC1412892) and counted using a Cellometer K2. Remaining cells were then pelleted at 400*g* for 5 minutes at 4°C and resuspended at 1×10^6^ cells/mL in ice-cold BamBanker medium (Bulldog Bio BB02). Cell suspensions were aliquoted at 2.5-5.0×10^5^ cells and frozen directly at −80°C until sorting.

### Cryorecovery and flow cytometry sorting of human microglia

Frozen human brain single-cell suspensions were thawed rapidly in a 37°C water bath with swirling and transferred to fresh 50-mL conical tubes. Cell suspensions were then diluted slowly with pre-warmed RPMI 1640 (Fisher MT10041CV) + 5% FBS (Gibco 26140079), 50µL every 5 seconds with agitation until 1.5mL, 100µL every 5 seconds with agitation until 5mL total volume. The resultant suspension was then filtered through a 70µm strainer and washed with 500µL RPMI + 5% FBS and spun down at 400*g* for 10 minutes at RT. Cells were then resuspended in 50µL ice-cold 1:10 human TruStain FcX (BioLegend 422302) in MACS buffer and incubated for at least 5 minutes on ice. A 2µL aliquot was then counted on a Cellometer K2 as above. Samples were then mixed with 50µL antibody cocktail / 1×10^6^ cells (minimum 50µL; see cocktail below) and incubated at 4°C in the dark for 30 minutes. Cells were then diluted with 900µL ice-cold MACS buffer and pelleted at 400*g* for 5 minutes at 4°C, and resuspended in 400µL ice-cold MACS buffer. Immediately before sorting, suspensions were filtered through a 70-µm filter, rinsed with 100-600µL MACS buffer. SYTOX stain was then added at 1µL and the suspension was mixed thoroughly. Cells were then sorted into PBS + 4% BSA (Millipore-Sigma A1595-50ML) in DNA lo-bind tubes (Eppendorf 022431005) using a FACS ARIA SORP in a BioProtect IV-LE-Bio Containment Hood with a 100µm nozzle. For scRNA-seq, cells were sorted as singlet, non-debris, live (SYTOX^-^), CD56^-^, CD15^-^, CD45^+^, HLA-DR^+^ events. Microglia were further subdivided as [CD14/CD3/CD19]^-^ events for flow cytometry.

### Single-cell RNA-seq

Sorted cells were diluted to 1.5mL with BamBanker medium and immediately pelleted at 400*g* for 5 minutes at 4°C. Cells were then resuspended in PBS + 4% BSA at ∼1×10^6^ cells/mL. Cell concentrations and viability were confirmed using a Cellometer K2 as above. Libraries were then generated using the 10X Genomics 5’ V2 kit, according to manufacturer’s instructions (CG000331 Rev A), using a 10X Genomics Chromium Controller. After quality checks, single-cell RNA-seq libraries were pooled and sequenced on a NovaSeq 6000 instrument using an S1 flow cell (Illumina 20028319).

### Single-cell RNA-seq analysis and processing

Data were processed using the Cell Ranger 7.0.1 pipeline (10x Genomics) with intronic reads disabled. To enable detection of viral RNA, reads were aligned to a custom hybrid genome containing GRCh38.93 and SARS-CoV-2 (NC_045512.2). An additional negative strand transcript spanning the entirety of the SARS-CoV-2 genome was then added to the GTF and GFF files to enable detection of SARS-CoV-2 replication as described in^7^. Samples were genetically demultiplexed using the supplied “common_variants_grch38.vcf” reference, stripped of genetically-defined doublet cells. Putative heterotypic doublets were then flagged for removal using scrublet 0.2.3 with manual thresholding of doublet scores, before removal using custom scripts in R^67^. Detection and removal of empty droplets was then performed using cellbender 0.2.0 using GPU optimization with a 40GB Tesla A100 GPU^68^. Where applicable, expected cell numbers were determined using the 10X Genomics 5’ V2 kit manual (CG000331 Rev A). Thresholding of initial filtering and preprocessing was performed using Seurat 4.2.1^69^, followed by integration using SCVI within SCVItools 0.14.3^70^, and re-imported into Seurat for all clustering, dimensional reduction, and all downstream high-level analysis using inbuilt Seurat functions and custom scripts in R. All manipulations in Seurat were performed with the aid of tidyseurat 0.5.3^71^. Normalization was performed using SCTransform 0.3.5^72^, and clustering was performed using the Leiden algorithm. Default parameters were otherwise used unless directly specified^73^.

### Pseudobulk differential expression analysis

Pseudobulk analysis was performed using the “bulkDEA” function in the “Seurat_pseudobulk_DEA.R” script in the NUPulmonary/utils repository. Briefly, raw counts (object@assays$RNA@counts) were aggregated by sum by sample (e.g. mouse or patient) and major cell type (e.g. microglia, CD8+ T cells) and passed from Seurat to DESeq2 1.34.0^74^ with relevant metadata. For human data differential expression analysis (DEA) was performed by group, i.e. COVID-19 vs control. Size factor estimation, dispersion fitting, and Wald tests were performed using the DESeq function in DESeq2. “Parametric” and “local” models of dispersion were compared visually for goodness-of-fit, and the most reasonable fit was chosen. Results were then extracted using the results function with alpha set to 0.05. For mouse data, DEA was performed across a combined factor of age and IAV treatment group, e.g. old_acute vs young_acute. Default parameters were used unless otherwise specified. In all plots of pseudobulk gene counts, p-values shown are FDR-corrected p-values directly from DESeq2 analysis. For gene-set-enrichment analysis (GSEA), the fgsea 1.20.0 package was used^75^. “Hallmark” gene set lists were downloaded from the Molecular Signatures Database (MSigDB) 7.5.1 at http://www.gsea-msigdb.org/gsea/downloads.jsp^29^. Enrichment analysis was then performed for all gene sets simultaneously using the “fgseaMultilevel” method using gene-level Wald statistics as rankings and default parameters.

### Multiplexed cytokine assays

In-house assays: Cytokine levels in matched BAL fluid and plasma collected from patients were measured using the multiplexed human cytokine/chemokine magnetic bead kits from Millipore (HCYTMAG-60K-PX41 and HCYP2MAG-62K) according to manufacturer’s protocol (HCYTOMAG-60K Rev. 18-MAY-2017). Briefly, frozen BAL and plasma samples were thawed, spun at 500rcf for 5 minutes to clarify, and 25uL of sample was added to 25uL of premixed magnetic beads (minus the beads for RANTES, PDGF-AA, and PDGF-AB/BB, which were excluded from analysis) in the provided 96 well plate, incubated at 4°C for 4 hours, washed, and then sequentially labeled with 25uL of detection antibodies followed by 25uL of streptavidin-phycoerythrin prior to analysis using a Luminex® 200 system. Raw MFI, bead counts, and standard concentrations were exported and analyzed as described below. CRO assays: Remaining assays were performed by Eve Technologies (Calgary, Alberta, Canada). Samples were thawed and aliquoted at 100µL, frozen and shipped to the CRO on dry ice. The Human Cytokine/Chemokine 71-Plex Discovery Assay (HD71) was then performed on each sample. Custom outputs containing raw MFI values, standard curve concentrations, and bead counts for processing as described below.

### Multiplexed cytokine assay processing and analysis

Processing and high-level analysis were performed using custom scripts in R 4.1.1, which are included in the GitHub repository for this publication. Raw MFI values, beads counts, and standard concentrations were first stripped from the data output from either Exponent (in-house assays; Luminex) or bespoke output from Eve Technologies (Calgary, Alberta, Canada). MFI measurements with fewer than 50 bead counts were discarded. Standard curves for each cytokine were then fit for each assay run using self-starting 5-parameter logistic (5PL) models using drc 3.2-0^76^. Cutoffs for curves with low predictive value were then determined empirically using histograms MFI values vs standard concentrations to identify a bimodal distribution cutoff. For in-house assays, all values calculated using standard curves with MFI < 50 at 100pg/mL were discarded. For Eve Technologies assays, all values calculated using standard curves with MFI < 50 at 10pg/mL were discarded. Experimental values for each cytokine were then predicted using the ED function in drc with “absolute” value prediction. In rare cases where a 5PL model could not be hit for an individual cytokine-assay combination, these values were excluded (see Fig. S3). Values below the lower asymptote of the model were set to a concentration of 0pg/mL. Values above the upper asymptote were set to the value of the upper asymptote. Technical replicates (including those across assays) were collapsed by mean with NA values excluded. Plots of standard curves for each analyte for each assay were automatically exported and are available upon request. Analytes showing poor dynamic range, e.g. TNF-ɑ for all plasma samples, were excluded from further analysis. For calculations of cumulative exposure (AUC) by ICU day, a piecewise linear function was first fit for each patient for each analyte for all measurements during the patient’s stay using the approxfun function in R stats 4.1.1 using the “linear” method with n = 100. Initial measurements were carried out an additional day to represent the time of admission to first measurement, and final measurements were carried out an additional day to represent time until discharge or cessation of measurement. In rare cases where an initial measurement was missing, values were imputed by setting the initial measurement (day = 0) to the first measurement for the analyte/patient pair. Geometric integration was then performed using the integrate function from R stats 4.1.1 using the day of first measurement as the lower bound and the final day of measurement + 1 as the upper bound. Default parameters were used unless otherwise specified.

### Bulk cytokine deconvolution

Raw scRNA-seq counts from Grant et al. (GSE155249)^7^ were imported as an H5AD object using SCANPY 1.9.1^77^. Genes were then filtered to the union between genes detected by scRNA-seq (counts > 0) and analytes analyzed in multiplexed cytokine assays. Expression of each analyte-encoding gene was then summarized by mean using pandas 1.5.1^78^ and numpy 1.23.4^79^ and exported as a CSV for further analysis in R 4.1.1 using the environment described herein. For correlations between protein expression by multiplexed cytokine assay and cell-type- and cell-state-specific scRNA-seq counts, exact Spearman correlations were performed using cor.test in R stats 4.1.1.

### Single-molecule fluorescence *in situ* hybridization and immunofluorescence

Fixed human postmortem frontal lobe sections were collected as described above and transferred to sterile-filtered 20% sucrose in PBS for 24-48 hours at 4°C until fully equilibrated. This procedure was then repeated with 10% sucrose + 50% Scigen Tissue-Plus O.C.T. Compound (Fisher 23-730-571). Equilibrated samples were then embedded in 100% Scigen Tissue-Plus O.C.T. Compound and frozen on dry ice before being stored at −80°C indefinitely. Tissue sectioning and pretreatment was performed according to manufacturer’s instructions using the “Fixed-frozen tissue sample preparation and pretreatment” protocol from the RNAscope Multiplex Fluorescent Reagent Kit v2 User manual (ACD 323100-USM/Rev Date: 02272019) and using the RNAscope H_2_O_2_ and Protease Reagents kit (ACD 322381) and RNA-Protein Co-Detection Ancillary Kit (ACD 323180). Tissues were sectioned using a Cryocut 1800 cryostat (Reichert Jung) at 14µm. Sections were transferred directly to RT Bond 380 microslides (Matsunami 0380W) at 2-3 sections per slide. Samples were boiled in ACD co-detection target retrieval buffer for 5 minutes at 95-102°C with occasional stirring on a hot plate. All downstream steps were performed according to the RNAscope® Multiplex Fluorescent v2 Assay combined with Immunofluorescence - Integrated Co-Detection Workflow (MK 51-150/Rev B/ Effective Date: 02/11/2021). For IBA1 staining, sections were stained with an anti-IBA1 antibody at a 1:50 dilution (Abcam ab178847) for 1-2 hours at RT followed by 3 washes in PBS + 0.1% Tween 20 (PBS-T; Sigma P7949-100ML). Primary antibody was then fixed to the tissue for 30 minutes in 4% PFA for 30 minutes at RT followed by 3 washes with PBS-T. Staining by smFISH and mounting were performed as described in the RNAscope 4-plex Ancillary Kit for Multiplex Fluorescent Reagent Kit v2 protocol (ACD 323120-TN/Rev A/Draft Date 12172019) using the RNAscope Multiplex Fluorescent Detection Reagents kit v2 (ACD 323110) and RNAscope 4-Plex Ancillary Kit for Multiplex Fluorescent Kit v2 (ACD 323120). Sections were stained with probes against *CCL2* (channel 1; ACD 423811) conjugated to Opal 690 (Akoya FP1497001KT), *CDKN1A* (channel 2; ACD 311401-C2) conjugated to Opal Polaris 780 (Akoya FP1501001KT), and *IL1B* (channel 4; ACD 310361-C4) conjugated to Opal 620 (Akoya FP1495001KT). Expression of IBA1 was then visualized using a secondary antibody against rabbit IgG conjugated to Alexa-Fluor 488 (Life Technologies A21206) at a 1:500 dilution in Co-Detection Antibody Diluent for 30 minutes at RT in the dark followed by 3 washes with PBS-T. Targets were stained in the order C1, C4, secondary antibody, C2 to enable staining with Opal Polaris 780. Prior to counterstaining with DAPI, lipofuscin autofluorescence was quenched by treatment with 1x TrueBlack stain (Biotium 23007) in 70% ethanol for 30 seconds followed by 3 washes in PBS. Samples were then counterstained with ACD DAPI solution for 1 minute followed by 1 wash in PBS. Sections were mounted with an excess of Prolong Gold mountant (Thermo-Fisher P36930) using VWR No. 1.5 coverglass (VWR 48393-195), cured overnight at RT in dark, and sealed with clear nail polish.

### RNAscope imaging and image processing

Slides from human brain sections were prepared as described above and imaged with a 60X Plan Apo oil immersion Objective (NA 1.4) on a Nikon ECLIPSE Ti2 wide-field inverted microscope equipped with a Photometrics Iris 15 camera at the Northwestern University Center for Advanced Microscopy & Nikon Imaging Center. DAPI, Alexa-Fluor 488, Opal 620, Opal 690, and Opal Polaris 780 were captured using a filter set for DAPI (Chroma 49000), EGFP (Chroma 49002), DSRed (Chroma 49005), Cy5 (Chroma 49006), and Cy7 (Nikon 96377), respectively, using LED illumination. Z-stacks were acquired for all images shown at 0.3µm per optical section. Image processing was then performed using a custom macro in FIJI/ImageJ version 2.9.0/1.54b^80,81^. Raw ND2 files were imported using the Bio-Formats plugin without rescaling and split by channel. Each channel was then flattened using a maximum Z projection. Background subtraction was then performed individually on each channel using the “Rolling Ball” algorithm individually for each channel using the “Subtract Background” function with a radius of 20 for all RNA-scope targets, 50 for DAPI/DNA, and 100 for IBA1. Individual channels were then rescaled with the same LUT to enhance contrast. Channels were then merged and pseudocolored using the “Merge Channels” function, converted into an RGB color TIFF, and exported. Individual channels were then inverted and exported as 16-bit TIFFs.

### Statistical analysis and data visualization

Statistical analysis was performed using R 4.1.1^82^ with tidyverse version 1.3.1^83^. For all comparisons, normality was first assessed using a Shapiro–Wilk test and manual examination of distributions. For parameters that exhibited a clear lack of normality, nonparametric tests were used. In cases of multiple testing, P values were corrected using FDR correction. Adjusted P values <0.05 were considered significant. Two-sided statistical tests were performed in all cases. Plotting was using ggplot 2.3.4.0 unless otherwise noted^84^. Comparisons for these figures were added using ggsignif 0.6.3^85^. Heat maps were generated using ComplexHeatmap 2.10.0 with clustering using Ward’s Method (D2) with Euclidean distance as the distance metric^86^. Figure layouts were generated using patchwork 1.1.2 and edited in Adobe Illustrator 2023. In all box plots, box limits represent the interquartile range (IQR) with a center line at the median. Whiskers represent the largest point within 1.5× IQR. All points are overlaid. Outlier points are included in these overlaid points but not shown explicitly.

## Tables

**Table S1.**
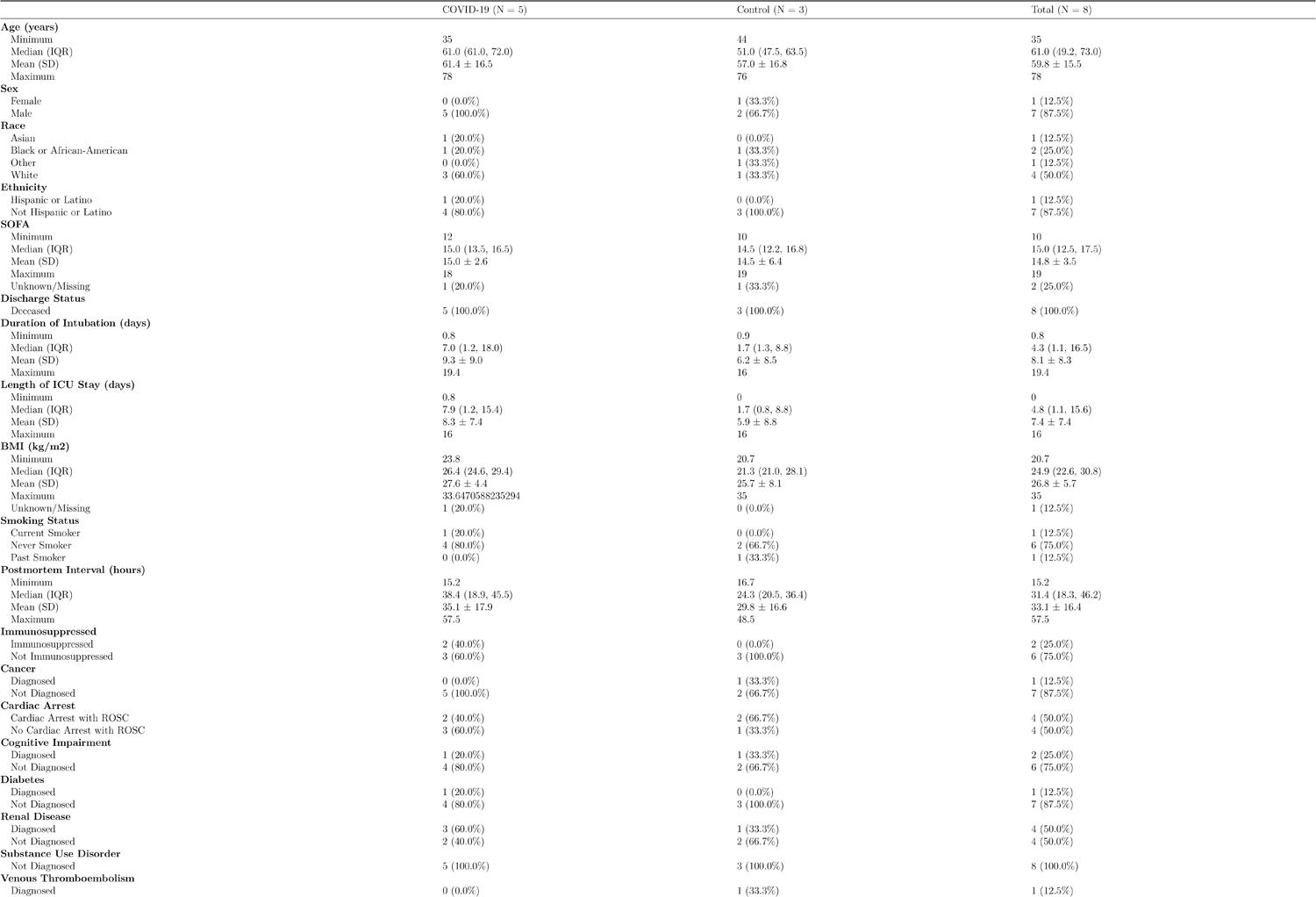
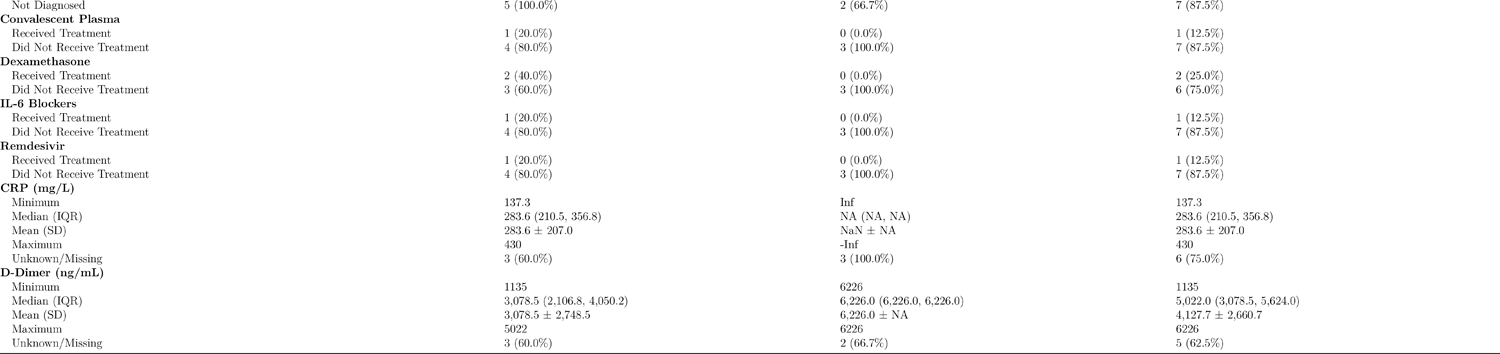
Demographics of the postmortem brain scRNA-seq cohort. Frontal lobe samples were collected in the course of autopsy at Northwestern Memorial Hospital.

**Table S2.**
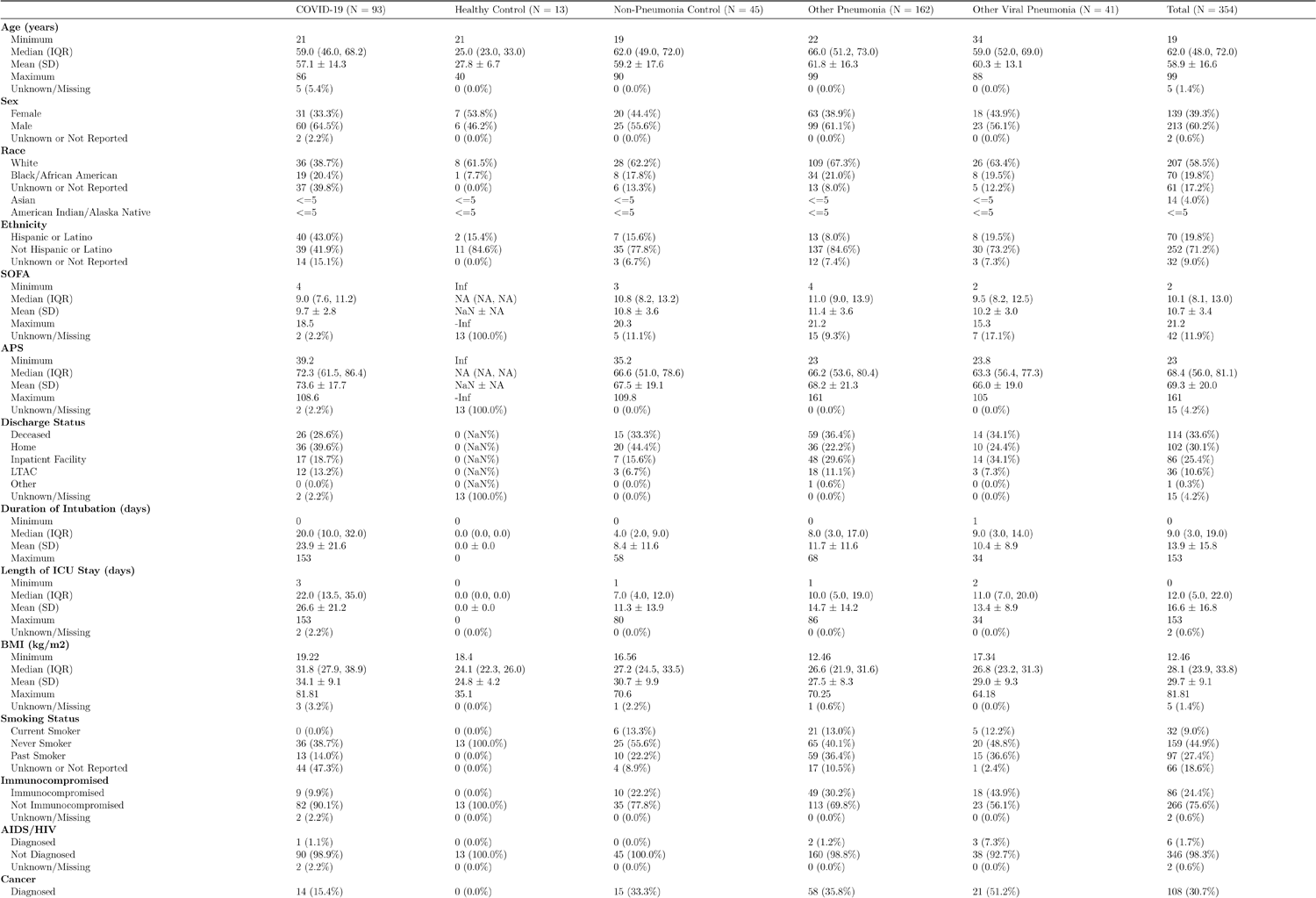
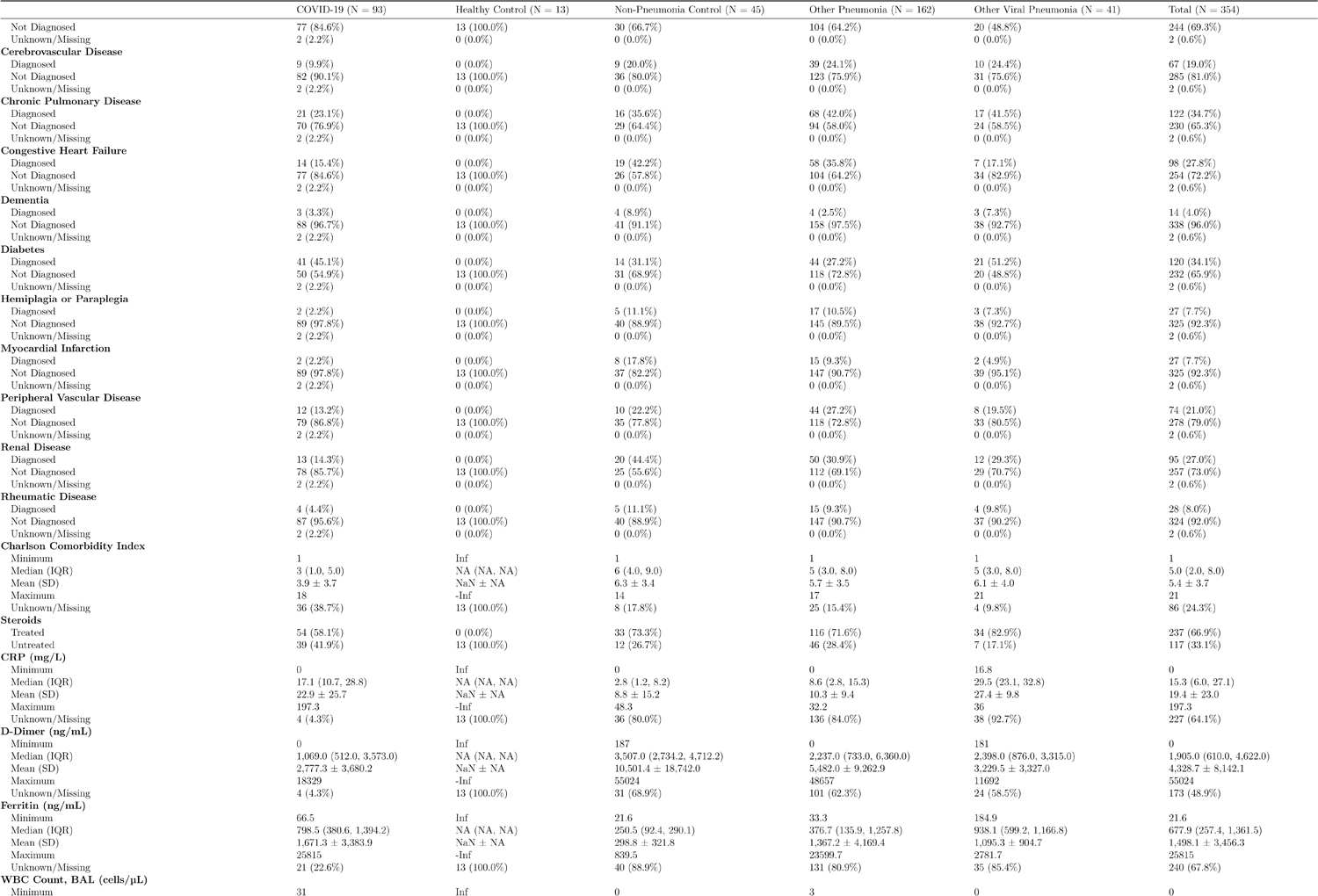
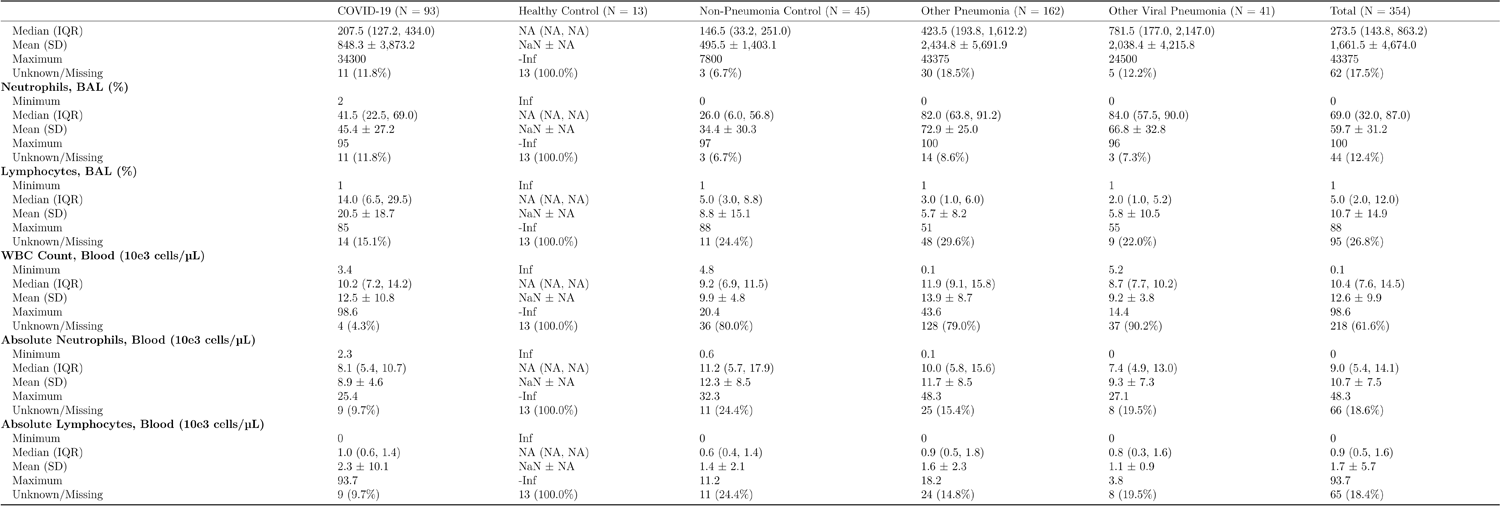
Demographics of the SCRIPT cytokines cohort.

**Table S3.** List of reagents used for FACS sorting of human neuroimmune cells.

| Laser | Filter | Dye | Antigen | Clone | Dilution | Catalog Number | RR ID |
| --- | --- | --- | --- | --- | --- | --- | --- |
| 305 | 750/50 | BUV737 | CD56 | NCAM16.2 | 1:10 | BD 612766 | AB_2813880 |
| 405 | 450/50 | eFluor450 | HLA-DR | L243 | 1:20 | Thermo-Fisher 48-9952-42 | AB_1603291 |
| 552 | 575/25 | PE | CD14 | MΦP9 | 1:10 | BD 562691 | AB_2737725 |
| 552 | 575/25 | PE | CD3 | SK7 | 1:20 | Thermo-Fisher 12-0036-42 | AB_10805512 |
| 552 | 575/25 | PE | CD19 | J3-119 | 1:10 | Beckman Coulter IM1285U | AB_10640419 |
| 488 | 530/30 | SYTOX Green |  |  | 1:500-1:1000 | Thermo-Fisher S7020 |  |
| 640 | 670/39 | APC | CD15 | W6D3 | 1:10 | BioLegend 323008 | AB_756014 |
| 640 | 780/60 | APC-Cy7 | CD45 | HI30 | 1:10 | CST 77322 | AB_2799894 |

